# Unique intramolecular oxidative rearrangement N-nitrosation mechanism of non-heam iron-containing enzyme SznF

**DOI:** 10.1101/2020.12.21.423711

**Authors:** Junkai Wang, Xixi Wang, Qingwen Ouyang, Wei Liu, Hongwei Tan, Xichen Li, Guangju Chen

## Abstract

Non-heam iron-dependent enzyme SznF catalyzes a critical step of the L-arginine derived guanidine group rearrangement to produce the N-nitrosourea pharmacophore in the process of SZN biosynthesis. The intramolecular oxidative rearrangement process is accomplished in the Fe(II)-containing active site located at the cupin domain of SznF, with which the catalytic mechanism remains elusive. In this work, density functional theory methods have been employed to investigate possible catalytic mechanisms of SznF. The N-nitrosation reaction in SznF was found to follow an energetically favorable pathway which includes six consecutive steps: (1) formation of Fe^II^-superoxo species with dioxgen binding on the iron center; (2) superoxo group attacking on the C^ε^ of substrate to form the peroxo-bridge complex; (3) C^ε^-N^ω^ bond homolysis to release N^ω^O; (4) peroxo bridge heterolytic cleavage; (5) deprotonation of 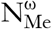 by Fe-O group; (6) the 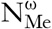 couples with the N^ω^O group and generates the N-nitroso product. The reaction proceeds in an unexpected way during which the electrons shuttle among two NO groups of the substrate and the peroxo moiety to promote C^ε^-N^ω^ bond homolysis and O-O bond heterolysis sequentially without generating high-valent Fe-O species, which is distinct from any known reactions catalyzed by the iron-containing enzyme. The unusual mechanism of SznF shed light on the area of enzymatic N-nitrosation reactions.

## Introduction

Streptozotocin (SZN) is a natural N-nitrosourea product derived from *Streptomyces achromogenes*,^1^ which is an approved chemotherapeutic agent for cancer with a commercial name Zanosar.^2^ For a long time, the biosynthetic pathway to produce N-nitrosourea pharmacophore of SZN remains unknown, until very recently, Ng et al. located a non-heam iron-dependent enzyme, SznF, which catalyzes a critical step of the L-arginine derived guanidine group rearrangement to produce the N-nitrosourea pharmacophore in the process of SZN biosynthesis.^3^ Structural characterization and mutagenesis of SznF indicated that it uses two separate active sites with different iron-containing metallocofactors to accomplish a stepwise transformation from N^ω^-methyl-L-arginine (L-NMA) to an N-nitrosourea product, of which the central domain catalyzes two successive N-hydroxylation reactions of L-NMA,^4^ while the cupin domain performs subsequent critical N-nitrosation on the N-hydroxylation intermediate N^δ^-hydroxy-N^ω^-hydroxy-N^ω^-methyl-L-arginine (**1**) to produce N^δ^-hydroxy-N^ω^-methyl-N^ω^-nitroso-citrulline (**2**) (Scheme 1a). Interestingly, this later reaction does not resemble any known cupin enzyme mediated transformation.^3^ It is also distinct from common N–N bond formation reactions in biosynthesis and chemical synthesis. Nitric oxide reductases (NORs) and flavo-diiron NO reductases (FNORs) are two identified enzymes among the relatively small group which get involved in natural N-N bond formation process.^5^ Both of them are able to reduce NO to N2O by means of external source electrons. NORs and FNORs are equipped with binuclear active center, by which two NO molecules are activated on one of the metallic sites respectively. Nevertheless, as a four-electron oxidation process fully coupled to the reduction of O2, the N-nitrosation reaction catalyzed by SznF is accomplished by a single metallic center without external reductant, which is very rare in both enzymology and organic synthesis. The elements of the N-N formation reaction in SznF resemble that of nitric oxide synthases (NOSs), which oxidizes the guanidine group of L-arginine to generate nitric oxide and L-citrulline,^6^ nevertheless SznF lacks the heam center of NOSs. Previously characterized in vivo N-nitrosation reaction routes employ external nitrite or NO^+^ as the active reaction species (Scheme 1b), while ^15^N isotope label experiment revealed that the two nitrogen atoms in the N-nitroso group of SZN are both from the same guanidyl moiety of arginine.^3^ Therefore, all the experimental evidence indicates that the N-nitrosation reaction catalyzed by SnzF follows a unique intramolecular oxidative rearrangement mechanism which remains elusive.

**Scheme 1.**
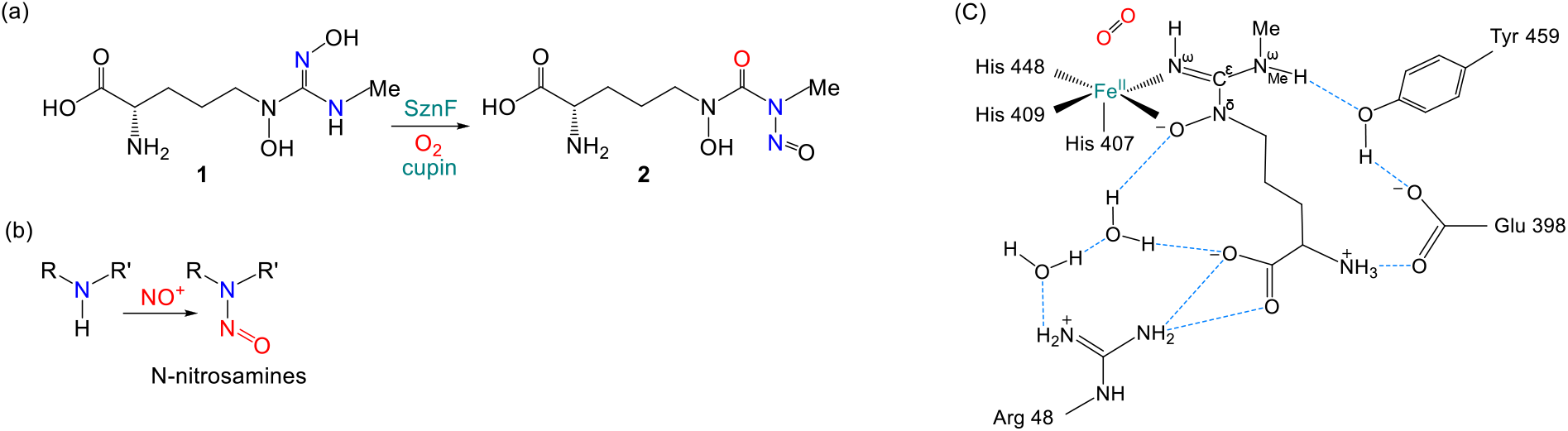
(a) SznF catalyzes the oxidative rearrangement of N^δ^-hydroxy-N^ω^-hydroxy-N^ω^-methyl-L-arginine (**1**) to generate the *N*-nitrosourea (N^δ^-hydroxy-N^ω^-methyl-N^ω^-nitroso-citrulline, **2**). (b) The common N-nitrosation reaction with external NO^+^. (c) Ligand-interaction map of the active site of SznF. The hydrogen bonds among the substrate analogue L-HMA, selected second shell residues and crystal water molecules are showed by blue dashed lines.

The X-ray structure of SznF complexed with the intermediate N^δ^-hydroxy-N^ω^-methyl-L-arginine (L-HMA) revealed the structural characteristics of its cupin domain (Scheme 1c). The active center features a 3-His (H407, H409 and H448) Fe^II^ binding triad, while the intermediate L-HMA acts as a bidentate ligand with N^δ^-hydroxyl group and one of its N^ω^ coordinating on the iron center, which leaves a vacancy site of iron for O2 binding. The methylated 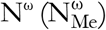 of L-HMA enrolls in a hydrogen bond chain through two residues Tyr459 and Glu398 to the ammonium terminal of the L-HMA, which stabilizes L-HMA in the active site along with the other hydrogen bond network formed among the carboxylate group of L-NMA, Arg48 and two crystalline water molecules (WAT713 and WAT828). The active center structure of SznF is very similar to that of 2-(oxo)-glutarate/Fe^II^-dependent dioxygenases (2-ODDs)^7^ and extradiol aromatic ring-cleaving dioxygenases (EDOs),^8^ implying potential mechanism similarity to them. Ng et al. proposed a possible catalytic mechanism for SznF (Path I, Scheme 2), which starts from the substrate **1** binding on the ferrous center in an orientation as that of L-HMA in the X-ray structure. Then, after O_2_ reduced to superoxo by an electron from the iron center, it adds on the guanidino-C (C^ε^) of the substrate, which propels the N^ω^O group ligating on the iron center to be broken off from the substrate and forms a cyclic peroxycitrulline intermediate (IntB1, Path I, Scheme 2). To complete this reaction step, the substrate needs to deprotonate on 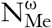 so that it is able to form a double bond with C^ε^. These proposed initial stages are very similar to the processes catalyzed by 2-ODDs that converts 2-(oxo)-glutarate to a ferryl-succinate complex. In the following O-O bond cleavage step, the iron center is oxidized to Fe^IV^=O complex (IntC1, Path I, Scheme 2). With C^ε^ of the substrate converting to a carbonyl, 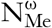 forms an amidate nucleophile, which then attacks the N^ω^O group coordinating on the iron and accomplishes N-nitrosation. Since the crossover experiment indicated that the nitrogen atoms in product **2** are from an intact guanidine group, which requires that the transient Fe^IV^=O complex intermediate does not exchange NO prior to N-N bond formation. Ng et al. therefore proposed an alternative mechanism (Path II, scheme 2) which avoids the formation of the Fe^IV^=O complex. In this second pathway, after the superoxo attacking the C^ε^, the N^ω^-C^ε^ bond in the substrate doesn’t break down but turns to form a diaziridine intermediate (IntC_2_, Path II, Scheme 2), which then experiences intramolecular rearrangement and O-O bond cleavage to generate the N-nitrosation product **2**. The proposal of this second pathway was inspired by EDOs. The most important difference between the two proposed pathways is the oxidation state of the iron center. In the Path I, the iron center converts to a high-valent ferryl state by donating two electrons to the peroxo moiety to promote O-O cleavage. While in Path II, after going through a transient Fe^III^-superoxo adduct, the iron center is reduced to ferrous state by the electron donated from the substrate. Though such an obvious difference between two mechanisms provides the opportunity to validate them through experiment, it is also intriguing to thoroughly investigate them with DFT calculations. For the catalytic mechanisms of SznF, there are also other interesting aspects that are worth studying. In the Path I, substrate **1** is proposed to be deprotonated on the 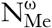 to facilitate the N-N formation after O-O bond cleavage. However, deprotonation of 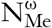 would alter the resonance structure of the substrate thus change its ligation to the iron center, which might result in an unexpected impact on the reaction. 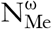 deprotonation is also strongly dependent on the protein environment around it. Therefore, the protonation state of 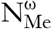 and its influence on the catalytic reaction remain elusive. In addition, in the proposed Path I, the C^ε^-N^ω^ bond breaks down prior to O-O cleavage after the superoxo added on the C^ε^. Whereas in the reactions catalyzed by the non-heam iron enzymes, such as protocatechuate 3,4-dioxygenase (3,4-PCD),^9^ *myo*-inositoloxygenase (MIOX),^10^ EDOs and *phn* operon (PhnZ),^11^ O-O bond usually cleaves immediately after peroxo bridge formation. Whether SnzF follows a similar reaction sequence to cleave the O-O bond firstly is still unclear. In 2-ODDs and EDOs, the iron center is coordinated by two histidines and a negative charge residue glutamate or aspartate, which enhances the reducibility of Fe^II^ in the enzymatic reactions. The iron center of SznF is coordinated by three neutral histidine residues, whether the iron center is able to be stabilized at the ferryl state during reaction progress is an interesting question to be explored.

**Scheme 2.**
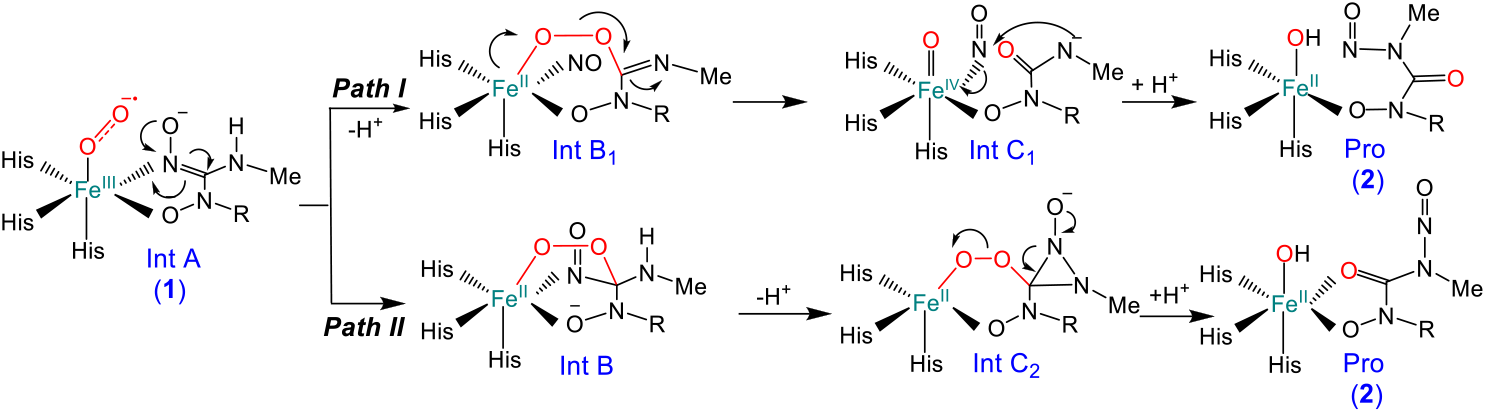
The two reaction pathways of SznF proposed by Ng et al.

In this study, extensive DFT calculations were performed to investigate the possible catalytic reaction mechanisms of SznF. Our calculation results revealed that the N-nitrosation reaction in SznF follows a unique mechanism that bypasses the formation of the high-valent Fe-O intermediate. Through a serial of electron rearrangements among two NO groups of the substrate and the O2 moiety, the reaction follows a pathway in which the C^ε^-N^ω^ bond cleaves followed by the O-O bond heterolysis. Then with the assistance from Fe^III^-O, the 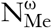 of the substrate deprotonates and completes N-nitrosation reaction through a nucleophilic reaction with the N^ω^O group. This unusual mechanism of SznF will reshape the current mechanistic understanding of enzymatic N-nitrosation reactions.

## Methods and Models

### Computational methods

For the iron-containing enzymes such as SznF, exchange interaction among the unpaired electrons plays important role in their activity and impacts the energetic processes of the catalytic reactions.^12^ Therefore, the magnitude of the energy barrier based on the DFT calculation is highly dependent on the exchange term embedded in the functional. A previous study on an iron-based enzyme AlkB indicated that hybrid functionals with different percentages of Hartree-Fock exchange gave out quantitatively different results, among which the functional ωB97XD has the best performance due to its ability to correctly reproduce the exchange interaction.^7a, 13^ Another important aspect that needs to concern by applying DFT calculations to the enzymatic reactions is whether the functional is capable of reproducing the dispersion interaction. The ubiquitous van der Waals interaction plays pivotal roles not only in the substrate binding process but also in the catalysis. In a study on the non-heam iron dependant enzyme intradiol dioxygenase, Borowski noted that the dispersion correction to the density functional is essential to accurately describe the dioxygen binding process in it.^14^ As a functional integrates empirical dispersion correction term, ωB97XD^15^ was therefore chosen to investigate the catalytic mechanism of SznF in this study. The geometric structures of all the stationary points, including reactant, intermediates and transition structures along the catalytic reaction pathways, were optimized by using the mixed basis sets, in which the LANL2TZ effective core potential^16^ was used for the Fe and the 6-31G(D, P) basis set^17^ for all the rest of the atoms. For each of the stationary points, broken-symmetry DFT approach was employed to investigate its different spin states. Analytic vibrational frequency calculations were performed on all the geometry structures to confirm that the correct stationary points were located and to obtain the corresponding zero-point energies (ZPE). Electronic energies were then evaluated with larger mixed basis sets by using the cc-pVTZ^18^ for all the atoms except LANL2TZ for Fe. Solvation effects were estimated by SMD continuum solvation model with dielectric constant set to 4 to mimic the protein environment.^19^

### Computational Model

To study the N-nitrosation mechanism catalyzed by SznF, a cluster model was built from the X-ray structure of SznF complexed with the substrate analogue L-HMA (PDB entry 6M9R). The iron center with the 3-His triad (His407, His409 and His448) was included in the cluster model. L-HMA was modified to the substrate N^δ^-hydroxy-N^ω^-hydroxyl-N^ω^-methyl-L-arginine (**1**) by adding a hydroxyl group on the N^ω^ which ligates on the iron center. An O_2_ molecule was added to the model to coordinate on the iron center. Three second shell residues, Tyr459, Glu398 and Arg48, as well as two crystalline water molecules, WAT713, and WAT828, which get involved in the hydrogen bond interactions with the substrate, were also built in the cluster model. The protonation states of all the residues were set based on the neutral pH environment. The substrate 1 was protonated on the ammonium terminal and deprotonated on the N^ω^, which has a net charge of −2. All the residues from SznF in the cluster models were truncated beyond alpha-carbon and saturated by hydrogen atoms. To maintain the overall structure of the active site, the boundary atoms were fixed in the optimization.

## Results and Discussion

### Binding of dioxygen to the iron center of SznF

DFT calculations were firstly performed to investigate the catalytic mechanisms of SznF proposed by Ng et al. The initial reactant (IntA, as shown in Scheme 2) is the complex formed by the substrate **1** and a dioxygen molecule binding on the iron center of SznF. Our calculation revealed that IntA is in a form of a ferrous-superoxo complex, in which the O2 moiety is reduced by an electron transferred from the substrate **1**, while the later one converts into a cationic radical ligand of the ferrous center. It is very similar to the situation of EDO from a previous study.^8, 20^ Whereas in 2-ODDs, its iron center instead of the substrate plays as the reductant in the dioxygen binding process due to its relative electron-rich ligand shell of iron which consists of a negatively charged glutamate and two histidine residues.^7b^ Depending on the number of unpaired electrons on the ferrous center and its coupling mode with the two single electrons on the superoxo and the substrate, IntA has three possible spin state, singlet (^1^IntA), triplet (^3^IntA) and quintet (^5^IntA), which further split into different electronic configurations depending on the coupling fashion among unpaired electrons. Extensive computational search for the electronic structures of IntA was therefore performed and corresponding results are summarized in Scheme 3. Our calculations indicated that the ground state of IntA is almost degenerated by the triplet (^3^IntA_HA_) and quintet (^5^IntA_HF_) (^5^IntA_HF_ is only 0.5 kcal/mol higher than ^3^IntA_HA_ in energy). While the singlet state lies 8.8 kcal/mol higher than the triplet. The difference between ^5^IntA_HF_ and ^3^IntA_HA_ is whether the unpaired electrons on the superoxo and the substrate cationic radical are ferromagnetically or antiferromagnetically coupled. Since our calculations indicated that the subsequent reaction step is dominated by the triplet state, we only discuss the triplet state of IntA (^3^IntA_HA_) here. Two possible coordination modes of superoxo on the iron center (end-on and side-on) were also examined (Scheme 4).^11, 20^ The calculation results indicated that in the singlet and triplet states of IntA, the superoxo can only adopt the end-on mode. Although we obtained side-on binding conformation of superoxo on the iron site in quintet state (Figure S1), which is energetically slightly higher (2.4 kcal/mol) than end-on ^5^IntA_HF_. Since the end-on binding conformation is feasible for the superoxo to attack on the C^ε^ in the subsequent reaction step, ^3^IntA_HA_ in which the superoxo binds on the iron center in end-on mode is therefore chosen as the initial reactant in this study. In the optimized ^3^IntA_HA_, the calculated spin population on the iron center is +3.8, corresponding to four α electrons on the ferrous center (Table 1). The gross spin population on the dioxygen and the N^ω^O moieties are −0.9 and −0.8 respectively, which, combined with the 1.32 Å O-O bond length, indicating a typical superoxo character of the dioxygen group in ^3^IntA_HA_ which is reduced by an α electron from the N^ω^O group of the substrate. As shown in Figure 1a, the Fe-O bond length is 2.01Å. And the dihedral angle of Fe-O-O-C^ε^ is measured as 38° with the distal oxygen atom in a distance of 3.27 Å to the C^ε^. Such an end-on conformation allows the distal oxygen to attack on the C^ε^ feasibly in the following reaction step. In ^3^IntA_HA_, by donating out an electron, the N^ω^O moiety merges in a delocalized conjugation bond formed among C^ε^, N^δ^, and 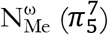, so that the N^ω^-O bond length is measured as 1.29 Å, which is 0.07 Å shorter than the N^δ^-O bond. And the five atoms, N^ω^O, C^ε^, N^δ^, and 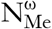 are in a coplanar fashion.

**Scheme 3.**
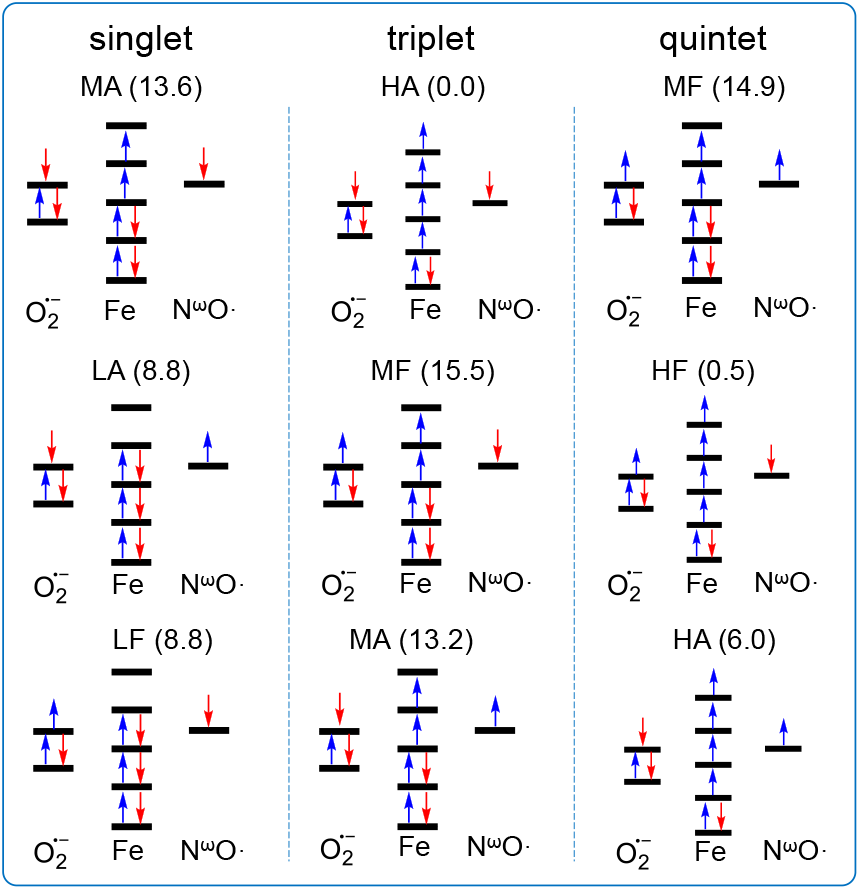
The electronic structures of the singlet, triplet and quintet states with the lowest energy (endon).

**Scheme 4.**
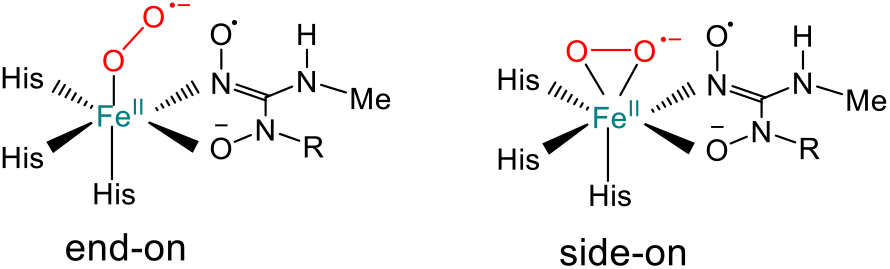
Different binding modes of superoxo on the iron center in IntA.

**Table 1.** The summarizations of spin populations on some important atoms for each of the structures.

| | $^3\text{IntA}_{\text{HA}}$ | $^3\text{TS}_{\text{AB}}$ | $^3\text{IntB}$ | $^3\text{TS}_{\text{BC}}$ | $^3\text{IntC}$ | $^3\text{TS}_{\text{CD}}$ | $^5\text{IntD}$ | $^5\text{TS}_{\text{DE}}$ | $^5\text{IntE}$ | $^5\text{TS}_{\text{EF}}$ | $^5\text{ProF}$ |
| --- | --- | --- | --- | --- | --- | --- | --- | --- | --- | --- | --- |
| Fe | +3.8 | +3.7 | +3.7 | +3.4 | +3.7 | +3.7 | +4.0 | +4.1 | +3.7 | +3.7 | +3.8 |
| O<sub>proximal</sub> | -0.3 | -0.2 | 0.0 | 0.0 | 0.0 | 0.0 | +0.7 | +0.4 | +0.1 | +0.1 | +0.1 |
| O<sub>distal</sub> | -0.6 | -0.4 | 0.0 | 0.0 | 0.0 | 0.0 | 0.0 | 0.0 | 0.0 | 0.0 | 0.0 |
| N<sup>ω</sup> | -0.3 | -0.4 | -0.5 | -0.7 | -1.2 | -1.0 | -0.6 | -0.6 | -0.6 | -0.5 | 0.0 |
| O (N<sup>ω</sup>) | -0.5 | -0.5 | -0.4 | -0.6 | -0.6 | -0.6 | -0.4 | -0.3 | -0.4 | -0.4 | 0.0 |
| N<sup>δ</sup> | -0.2 | -0.3 | -0.6 | 0.0 | 0.0 | 0.0 | 0.0 | 0.0 | +0.6 | +0.5 | 0.0 |
| O (N<sup>δ</sup>) | 0.0 | -0.1 | -0.4 | 0.0 | 0.0 | 0.0 | +0.1 | +0.2 | +0.4 | +0.4 | 0.0 |

**Figure 1.**
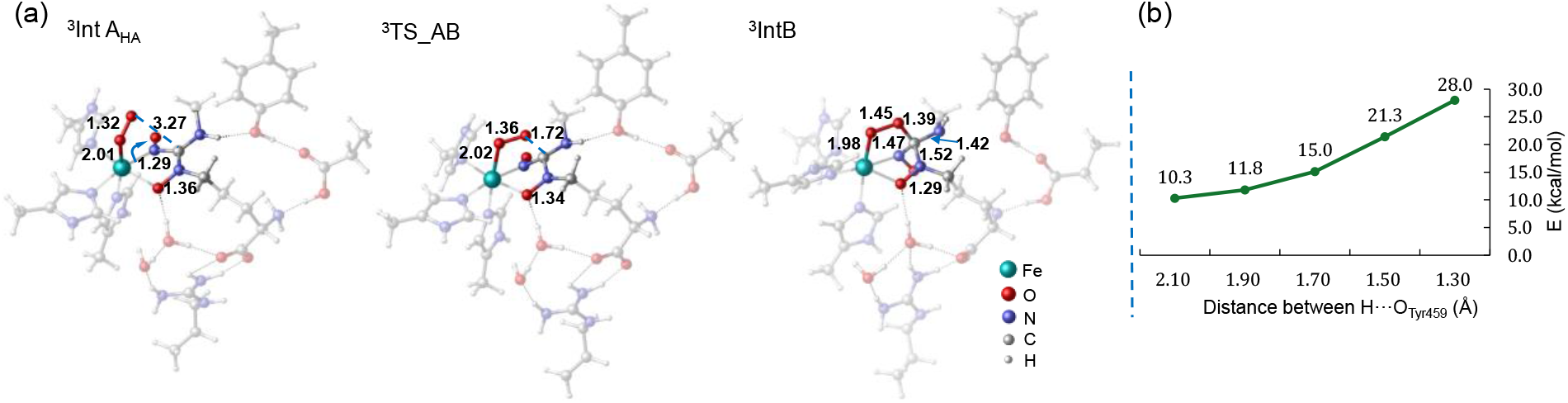
(a) The geometry structure of ^3^IntAHA, which is formed after dioxygen binding. The transition structure (^3^TS_AB) of the superoxo attacking substrate step and the formed peroxo-bridging intermediate (^3^IntB). Some key distances of the structures are labeled in the unit of Å. (b) The energy variations along the coordination of 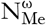 deprotonation by the Tyr459, which is obtained from the scan calculation with the step size of 0.2 Å.

### Superoxo addition on the substrate

In ^3^IntA_HA_, the superoxo is prepared as a radical with one β electron on it, whereas the substrate still needs to experience a complex electron rearrangement so that the distal oxygen of superoxo moiety attacks on the C^ε^ (Scheme S1). As shown in Figure 1a, from ^3^IntA_HA_, the N^ω^O moiety is polarized by the positive charge on the iron center. It triggers the following electron rearrangement in the substrate molecule: 1) the π bond between C^ε^ and N^ω^ breaks down and the two bond electrons become the lone pair electrons on N^ω^, 2) A single α electron is transferred from the N^δ^O moiety to C^ε^. The electron rearrangement promotes the C^ε^ to couple with the distal oxygen of superoxo and forms the cyclic peroxyarginine intermediate ^3^IntB. Our calculations indicated that this reaction step is the rate-limiting one of the whole catalytic processes. It costs 22.3 kcal/mol from ^3^IntA_HA_ to reach the optimal transition state ^3^TS_AB. The energy profile is shown in Figure 2. Though there is another possibility that the iron center plays as the reductant to donate electron to the O_2_ moiety in this step to generate a ferric state IntB, broken-symmetry DFT calculations indicated that it is relatively energetically unfavorable than the one with electron transferred from the N^δ^O (Figure S2). Since that the N^ω^O and N^δ^O moieties have very similar occupied frontier π^*^ orbitals which are close in their energy level, and the N^ω^O group has already shown that it has a higher potential than the iron center to donate out electron in the step dioxygen binding step, it is not strange that the N^δ^O instead of the iron center continues to supply the second electron to reduce the peroxo in this step. The optimized geometry structure of ^3^TS_AB is shown in Figure 1a. In ^3^TS_AB, the O-O bond elongates to 1.36 Å, with the distal O approaching C^ε^ in a distance of 1.72 Å. The tetrahedral conformation of C^ε^ reflects that it is converting from *sp*^2^ to *sp*^3^ hybridization. The three C^ε^-N bonds all stretch longer in ^3^TS_AB compared to that in ^3^IntA_HA_. The N^δ^-O bond, on the contrary, shortens to 1.34 Å, indicating that it is donating out electron from its π^*^ orbital. Spin population analysis of ^3^TS_AB confirms that an α electron is partially transferred from the N^δ^O group to the superoxo (Scheme S1). The down spin population summed on two oxygen atoms increases to −0.6, while that on N^δ^O decreases to −0.4 (Table 1). By crossing over the transition state ^3^TS_AB, it forms the peroxyarginine intermediate ^3^IntB. In ^3^IntB, the iron center remains in high spin ferrous state as that in ^3^IntA, while there are two unpaired electrons located on the N^δ^O and N^ω^O groups respectively, which antiferromagnetically couple with the four α electrons on the iron center. As shown in Figure 1a, the formation of C^ε^-O bond induces electronic and structural rearrangement in the substrate. In ^3^IntB, O-O bond length elongates to 1.45 Å. With C^ε^ converting to *sp*^3^ hybridization form, the conjugation bond between C^ε^ and N^ω^O group is therefore broken, which results in stretched C^ε^-N^ω^ bond length of 1.47 Å. The 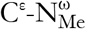 and C^ε^-N^δ^ bonds also elongate to 1.42 Å and 1.52 Å correspondingly. The observed excessively stretching of the C^ε^-N^δ^ bond obviously should be owed to that the N^δ^-O group loses an electron in this reaction and generates an electron hole on the N^δ^, which extremely polarizes the C^ε^-N^δ^ bond. On the contrary, losing an α electron from its anti-bonding π orbital results in strengthening of N^δ^-O bond, which shortens to 1.29 Å in ^3^IntB. The calculated gross spin population summed on the N^δ^-O group is −1.0. As proposed by Ng et al., from the peroxyarginine intermediate ^3^IntB, to follow the Path I, the substrate needs to deprotonate on 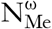 with the assistance of the two second shell residues Tyr459 and Glu398. However, our calculation revealed that it is unlikely for the 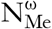 to deprotonate during this reaction step. First of all, 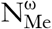 remains to be protonated in the optimized ^3^IntA and ^3^IntB. Moreover, We performed scan calculation along the coordinate with the proton moving from 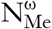 to the hydroxyl oxygen of Tyr459, which indicated that deprotonation of 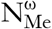 by Tyr459 in ^3^IntB is an energetically monotony increasing process as shown in Figure 1b. Furthermore, with the distance between H and O_Tyr459_ being shortened under 1.00 Å, 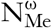 turns to abstract another proton from the ammonium group of the substrate.

**Figure 2.**
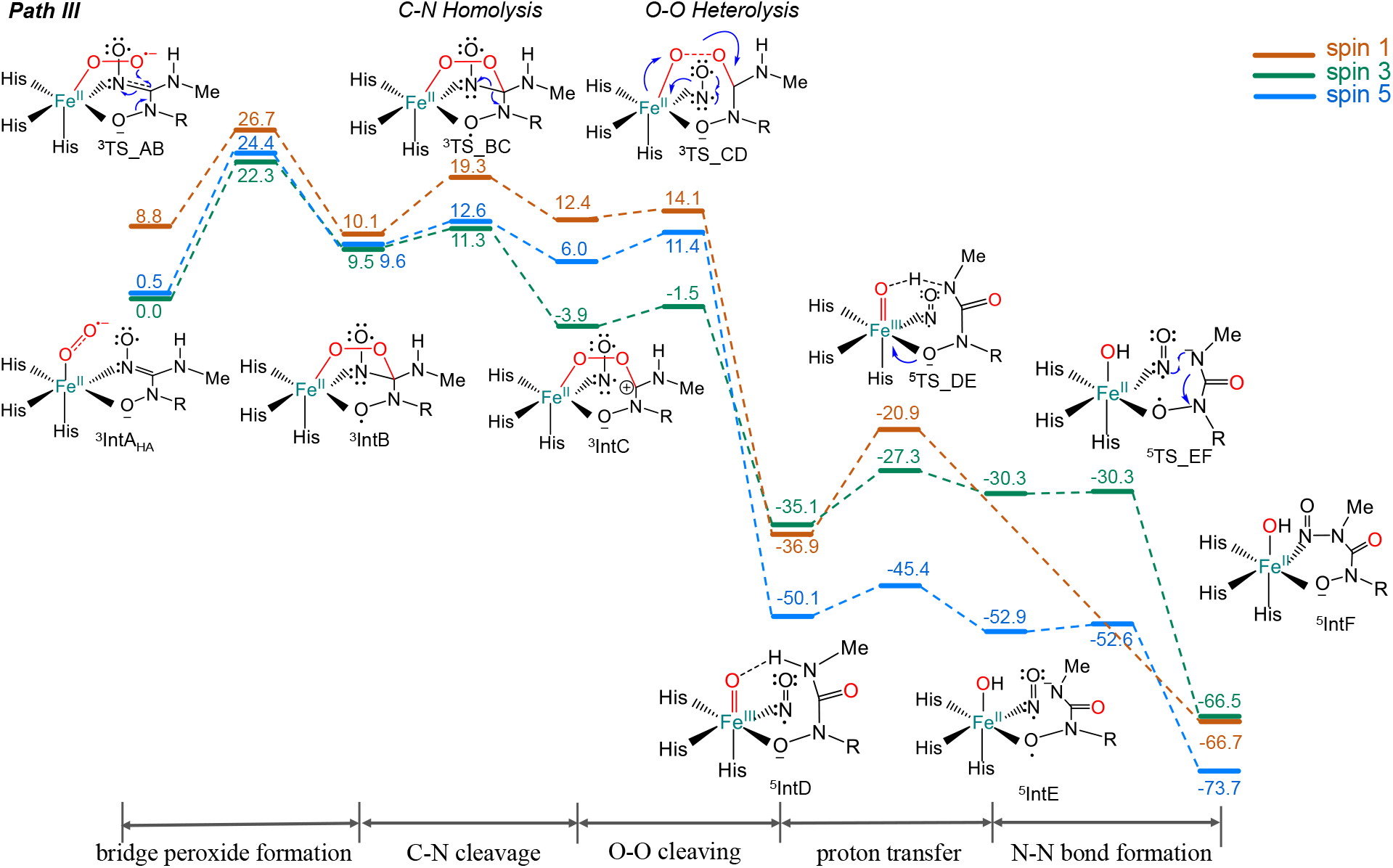
Free energy profile of the N-nitrosation mechanism catalyzed by SznF through intramolecular oxidative rearrangement. The relative energies are labeled in kcal/mol.

### N^ω^-C^ε^ Bond Cleavage

Though 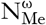 stays being protonated in ^3^IntB, our calculation indicated that N^ω^-C^ε^ bond cleavage could still proceed feasibly, except that in this step C^ε^ transfers an electron to the N^ω^O moiety but not to 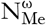, which leads to the intermediate ^3^IntC (shown in Scheme 5, Path III) by crossing over the transition state ^3^TS_BC with a very low energy barrier of 1.8 kcal/mol. In the optimized ^3^TS_BC, the N^ω^-C^ε^ bond length extends to 1.59 Å, while the N^ω^-O bond shortens to 1.24 Å (Figure 3). As evidenced by the calculated spin population, N^ω^-C^ε^ cleavage proceeds via a homolytic fashion. The spin population summed on the N^ω^O group decreases to −1.3 in ^3^TS_BC and further to −1.8 in the generated intermediate ^3^IntC, while the down spin population on the N^δ^O group is quenched in ^3^TS_BC and ^3^IntC, which confirms that a single α electron from the broken N^ω^-C^ε^ bond is transferred to the N^δ^O group, while the β electron goes to the N^ω^O group in this reaction step. In the ^3^IntC, C^ε^ converts to a carbocation and restores to planar *sp*^2^ conformation (Figure 3). The released N^ω^O^-^ is in triplet state and ligates on the iron center with Fe-N^ω^ distance of 2.01 Å. A noticeable structural characteristic of ^3^IntC is that the 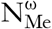 remains to be protonated with the 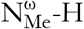 length of 1.01 Å, although the 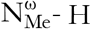 group has a hydrogen bond with Tyr459. The scan calculation also identified that deprotonation of 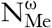 by Tyr459 in ^3^IntC is energetically unfavorable (Figure S5). Due to that deprotonation of 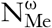 cannot be accomplished at this stage, the atom C^ε^ has to stay in the form of a carbocation in ^3^IntC. This C^ε^ carbocation polarizes the O-O bond adjacent to it and initiates the electron transfer cascade from the N^ω^O^-^ through the iron center and the O-O bond to the C^ε^, which finally leads to cleavage of the O-O bond (Scheme S1).

**Scheme 5.**
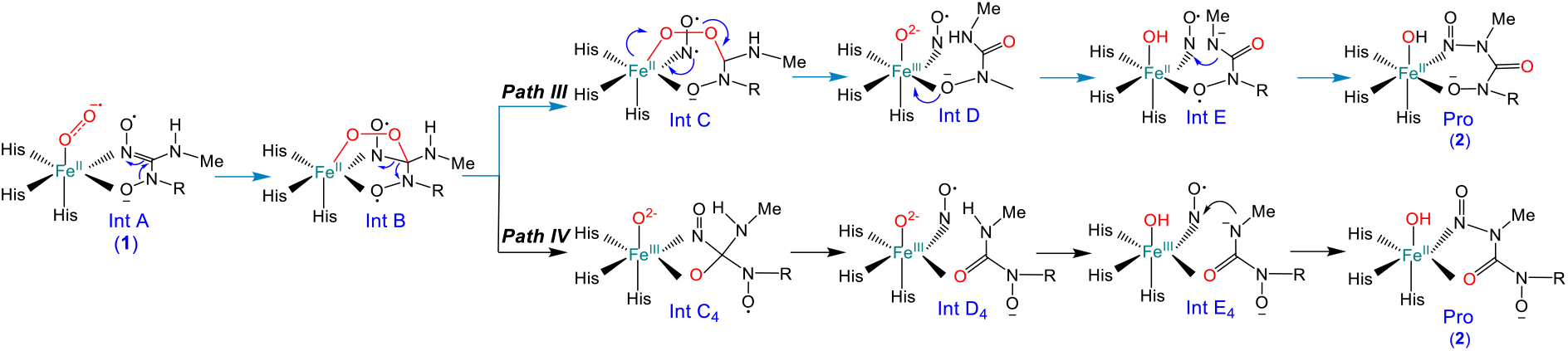
Two alternative possible pathways proposed in this work (Path III and Path IV). In Path III and IV, 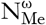 is deprotonated by Fe-O moiety after O-O bond cleavage. From peroxo-bridging intermediate Int B, by following Path III, N^ω^-C^ε^ bond cleaves prior to O-O bond cleavage. In the contrary, O-O bond cleaves prior to N^ω^-C^ε^ bond broken in Path IV.

**Figure 3.**
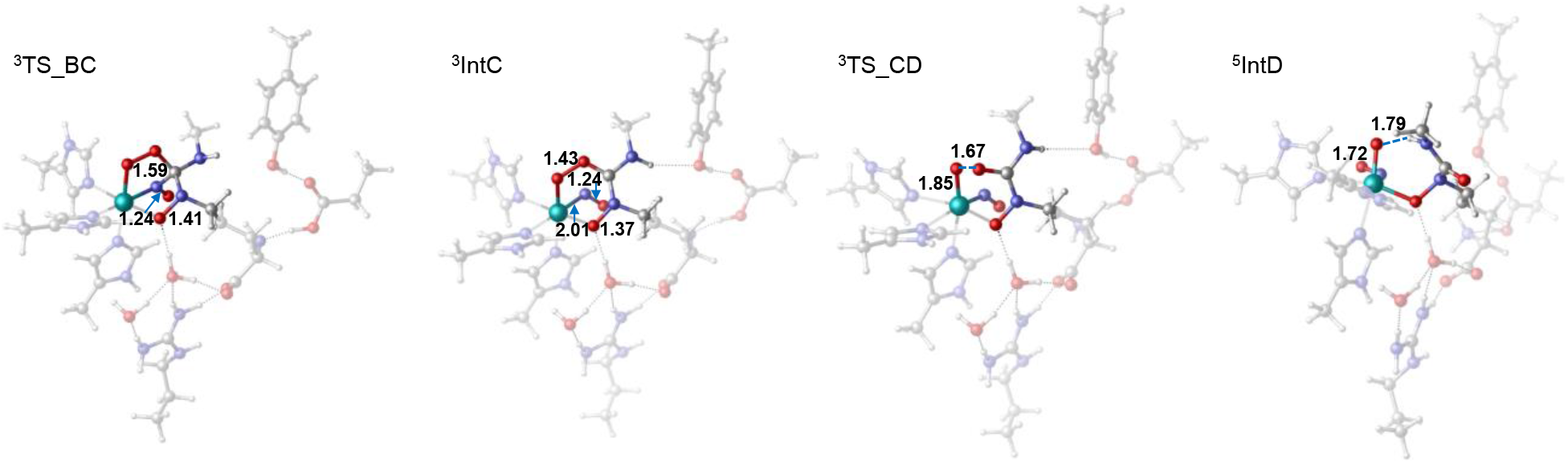
The structure of C-N homolytic cleavage transition state (^3^TS_BC) and the formed carbocation intermediate (^3^IntC). ^3^TS_CD is the geometry structures of O-O heterolytic cleavage transition state and ^5^IntD is the formed quintet state intermediate. The related distances (Å) are labeled in the structures.

### O-O Bond Cleavage

The O-O bond cleavage transition state ^3^TS_CD is only 2.4 kcal/mol higher than ^3^IntC, in which the O-O bond length extends to 1.67 Å with an imaginary frequency of 783 cm^-1^ (Figure 3). The transition state ^3^TS_CD is essentially one of a heterolytic cleavage, in which the spin populations on both of the two oxygen atoms are zero. From ^3^IntC, the O-O bond cleavage is exothermic by 46.2 kcal/mol and generates the intermediate IntD. The low energy barrier and extreme exothermicity of this O-O bond cleavage step results in that the subsequent reaction steps proceed very fast so that the released N^ω^O^-^ can hardly be exchanged during the reaction, which is consistent with the experimental observed results that the two nitrogen atoms in the N-nitroso group of SZN are both from the same guanidyl moiety of arginine. From ^3^IntC to IntD, the iron center and the N^ω^O^-^ anion each give out an electron in α and β spin to promote O-O bond cleavage. In the meantime, the iron center converts into a high spin ferric state, so that the reaction complex IntD experiences spin crossover and shifts to the quintet ground state. ^5^IntD is energetically 15.0 kcal/mol favorable than ^3^IntD. To the best of our knowledge, such an O-O bond cleavage process in SznF is distinct from that have been observed in most of the non-heam iron enzymes, in which the electron source to promote O-O bond cleavage usually is the iron center and generates Fe^IV^=O species after O-O bond heterolysis. It doesn’t resemble a recently studied diiron enzyme PhnZ either, by which the catalytic reaction goes through an unusual “inverse” heterolytic O–O cleavage transition state by abstracting two electrons from the substrate.^11, 14^

Spin population analysis of ^5^IntD revealed that the N^ω^O is a neutral molecule with a single β electron on it, which is antiferromagnetically coupled by the five α electrons populated on the Fe-O moiety. Though the calculated spin on the iron center and oxygen are +4.0 and +0.7 respectively, the 1.72 Å Fe-O bond length indicates that it is a Fe^III^=O species. The non-zero α spin on the oxide should be delocalized from the ferric center. It is interesting to notice that in the optimized ^5^IntD, the substrate is twisted around the C^ε^-N^δ^ bond after O-O bond cleavage and results in that the 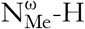 approaches the Fe^III^=O group to form a H-bond with it (Figure 3). The H…O distance is measured of 1.79 Å. It infers that deprotonation of 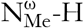 might be able to proceed at the stage of ^5^IntD with the assistance of Fe^III^=O. We therefore examined this deprotonation process with DFT calculations. The located 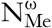 deprotonation transition state (^5^TS_DE) by the Fe^III^=O moiety is only 4.7 kcal/mol higher than ^5^IntD, which obviously could proceed readily at this stage. Interestingly, we found that deprotonation of 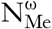 is accompanied by an electron transferred from the N^δ^O moiety to the iron center, which results in a serial of geometric changes in ^5^IntE as shown in Figure 4. 1) Fe^II^-OH^-^ elongates to 1.88 Å. 2) By losing an electron, there forms a 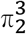 conjugation bond in N^δ^O group so that the atom N^δ^ converts from *sp*^3^ tetrahedral to planar conformation. 3) Correspondingly, N^δ^-O bond shortens to 1.29 Å from that of 1.43 Å in ^5^IntD. 4) The conformation alteration of N^δ^ enables the deprotonated 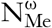 to twist and approach the N^ω^O ligand which is polarized by the iron center. The 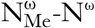 distance is 1.99 Å in ^5^IntE. Such a conformation is well prepared for the final nitrosation on 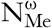.

**Figure 4.**
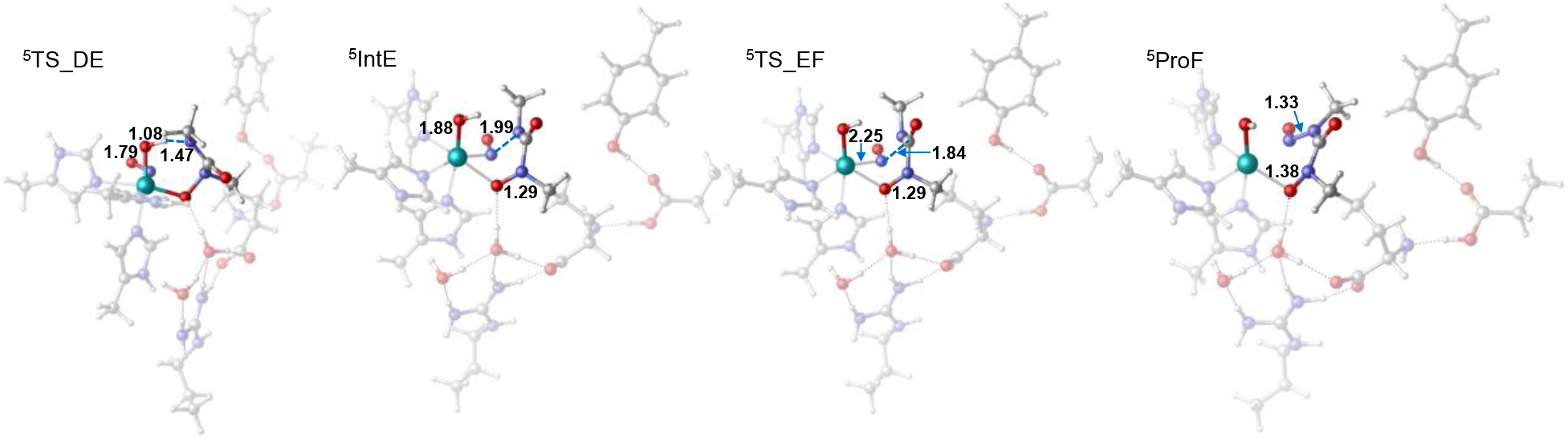
The transition structure of proton transfer step (^5^TS_DE) and the formed Fe^III^-O intermediate (^5^IntD). The ^5^TS_EF is the transition structure of the deprotonated 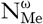 nucleophilic attacking the N^ω^O group which generates the N-nitrosourea product (^5^ProF). The key distances are labeled in the unit of Å.

### Formation of N-nitroso Group

From ^5^IntE, the N-N coupling step only needs to cross over a very low local energy barrier of 0.3 kcal/mol (almost barrierless) to reach the final product (^5^ProF). In the optimized ^5^TS_EF, the 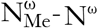 distance shortens to 1.84 Å, while N^ω^ detaches from the iron center with N^ω^-Fe distance stretching to 2.25 Å (Figure 3). The spin population of ^5^TS_EF is almost similar to that of ^5^IntE, indicating it is a nucleophile attacking transition state. Nucleophile coupling between 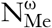 and N^ω^ triggers an electron rearrangement in which the single β electron on the N^ω^-O moiety transfers to the N^δ^O group and pairs with the single α electron on it. The electron rearrangement generates the N-nitrosation product ^5^PorF (N^δ^-hydroxy-N^ω^-methyl-N^ω^-nitroso-citrulline) and completes the catalytic cycle. This last step is exothermic for 20.8 kcal/mol.

### O-O Cleavage before C-N (Path IV)

From the peroxyarginine intermediate ^3^IntB, by following Path III catalyzed by SznF, C^ε^-N^ω^ bond cleaves and N^ω^O moiety is released as an anion ligand of the iron center. Nevertheless, previous investigation on EDO, diiron oxygenase PhnZ and MIOX revealed that the peroxo-bridging intermediates could take an alternative pathway in which the O-O bond of peroxo bridge cleaves preferentially.^8, 10–11^ It thus inspired us to explore the possible O-O bond cleavage process of ^3^IntB (Scheme 5, Path IV). Our calculations indicated that O-O bond cleavage pathway still proceeds on the optimal triplet state energy surface. The located transition state ^3^TS_BC4 is featured with homolysis characteristics based on the +0.3 and −0.4 spin on two oxygen atoms respectively (Figure S8). The energy barrier from ^3^IntB to reach ^3^TS_BC4 is 15.5 kcal/mol. ^3^TS_BC4 is 13.7 kcal/mol higher than the C^ε^-N^ω^ cleavage transition state ^3^TS_BC and 2.7 kcal/mol higher than the rate-limiting transition state ^3^TS_AB. Such a high energy barrier therefore abolishes the O-O cleavage pathway from the peroxyarginine intermediate ^3^IntB, though the following C^ε^-N^ω^ cleavage and N-nitrosation steps in this pathway are energetically feasible. The calculated energy barriers for the O-O cleavage step in PhnZ and EDO are both lower than that in SznF.^8, 11^ One possible reason is that SznF has a relative hydrophobic active site which prevents the peroxo bridge from protonation, while in PhnZ and EDO, the peroxo moiety is protonated before O-O bond cleavage. In addition, to promote O-O homolysis from ^3^IntB, a pair of electrons need to be transferred from the iron center and its ligand N^ω^O group to the peroxo moiety, which is also energetically unfavorable at this stage. On the contrary, by breaking down the C^ε^-N^ω^ bond of ^3^IntB, its two bond electrons are transferred to the N^ω^O and N^δ^O group respectively. The two NO groups are both polarized by their coordination bonds to the iron center so that the reaction only needs to cross over a very small energy barrier of 1.8 kcal/mol. Moreover, cleavage of C^ε^-N^ω^ bond could also facilitate the subsequent O-O bond broken. After C^ε^-N^ω^ bond homolysis, C^ε^ converts to a carbocation by transfer an electron to the N^δ^O group in ^3^IntC, which strongly polarizes its adjacent O-O bond thus promotes its heterolysis in the next reaction step. In the intermediate ^3^IntC after C^ε^-N^ω^ bond homolysis, the N^ω^O group is in a form of an anion, which also enhances its ability to assist O-O bond cleavage by donating an electron.

### Diaziridine Intermediate Pathway (Path II)

Ng et al. also proposed another possible N-nitrosation mechanism which goes through a diaziridine intermediate (IntC2, Path II, Scheme 1) from IntB.^3^ We further investigated this pathway with DFT calculations. Our calculations revealed that the reaction step to form the diaziridine intermediate is dominated by the quintet state. The only difference of ^5^IntB from ^3^IntB is that the single electron on the N^δ^O moiety is in α but not β spin, which is antiparallel to the β electron located on the N^ω^O group. Such an electronic arrangement enables N^δ^O moiety to accept the single β electron from N^ω^O so that the deprotonated 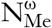 nucleophilic attacks on the N^ω^ to form the diaziridine intermediate. To follow this pathway, the substrate needs to deprotonate on 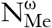, which is energetically highly unfavorable, hence it is not surprising that we found this pathway is actually forbidden in the enzymatic reaction by its extremely high energy barrier of 50.0 kcal/mol (Figure S11).

## Conclusion

In this study, DFT calculations were employed to study the reaction mechanism of the N-nitrosation reaction catalyzed by SznF. The energetic most favorable mechanism is divided into six steps:1) a dioxygen molecule binds to the active center and then is reduced by an electron transferred from the substrate to form the Fe^II^-superoxo complex; 2) superoxo group attacks on the C^ε^ of the substrate and forms a peroxyarginine intermediate; 3) C^ε^-N^ω^ bond homolytically cleaves and gives out a C^ε^ carbocation intermediate. 4) O-O bond heterolytic cleaves; 5) 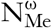 is deprotonated by Fe^III^-O group; (6) finally, the 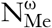 nucleophilic attacks the N^ω^O group and produces the N-nitroso group. The step in which the superoxo attacks on the substrate is the rate-determining step with 22.3 kcal/mol energy barrier. In contrast to the reactions proceeding in most of the iron-containing enzymes, the bridging peroxo intermediate in SznF doesn’t take the pathway to cleave O-O bond firstly. It homoytically cleaves C^ε^-N^ω^ bond and generates a C^ε^ carbocation intermediate after electron rearrangement. The catalytic reaction is featured with a very low energy barrier of the extreme exothermic O-O bond heterolysis step after C^ε^-N^ω^ bond cleavage, which results in that the reaction system hardly exchanges NO species prior to N-N bond formation. As a four-electron oxidation process, the most intriguing feature of this reaction is that during the whole catalytic cycle, the iron center switches between ferric and ferrous states but does not convert to a high-valent ferryl state.

## Supporting information

support information

