## Supplementary material for "Unique intramolecular oxidative rearrangement N-nitrosation mechanism of non-heam iron-containing enzyme SznF": support information: SI.pdf

**Contents**

Part I Supplementary of Path III. ........ S2
Part II The energy profile and O-O cleavage structures of Path IV. ........ S6
Part III The energy profile and structure of diaziridine intermediate in Path II. ........ S8

### Part I Supplementary of Path III.

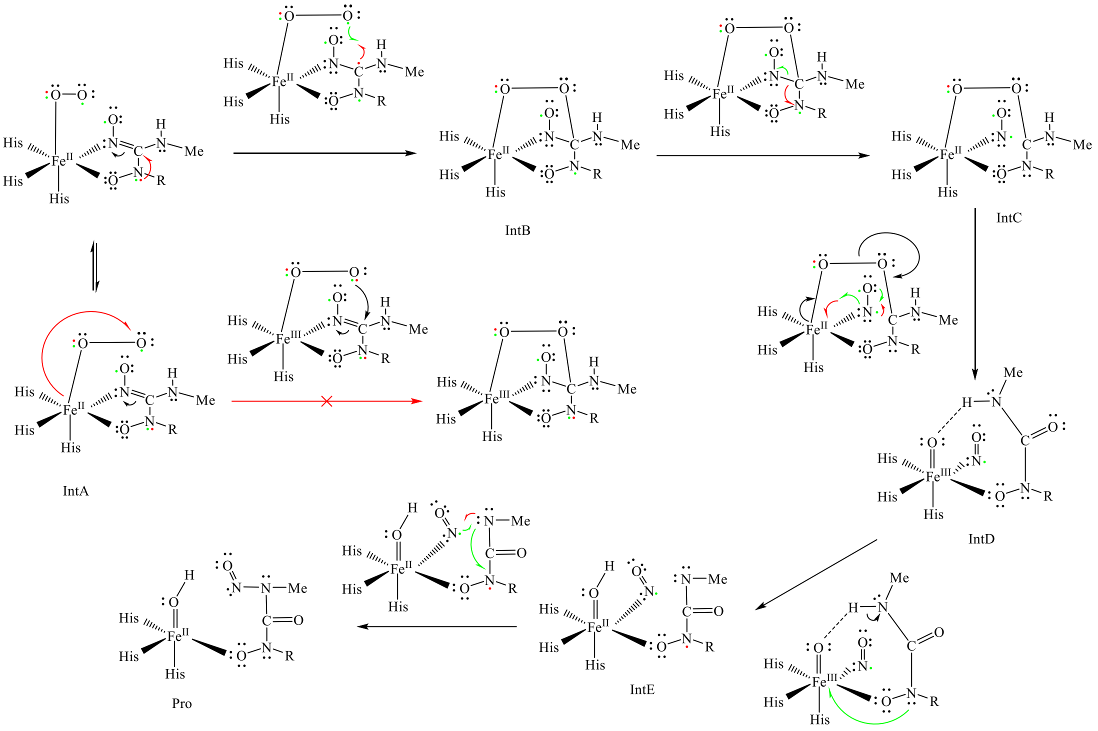


**Scheme S1.** The electronic structure changes along the Path III.

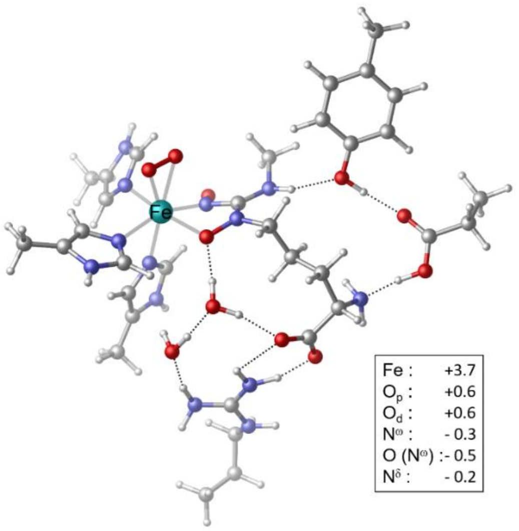


**Figure S1.** The structure of $^{5}$IntA$_{HF}$ with a side-on binding dioxygen group. The energy of this structure is 2.9 kcal/mol. The spin populations of some important atoms are summarized in the right corner.

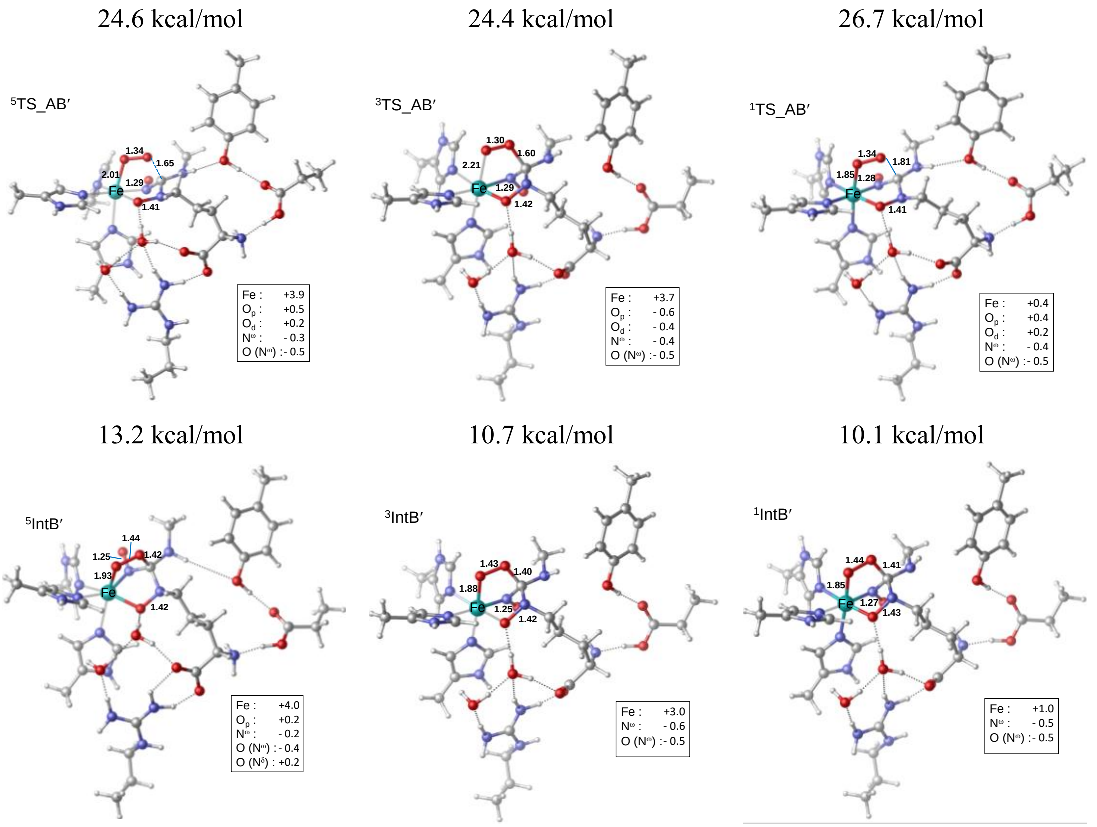


**Figure S2.** The structures of superoxo group attacking the $C^{\varepsilon}$ atom on the substrate, in which the electron of the iron center transfers to the dioxygen group to assistance the formation of $C^{\varepsilon}$-O bond. The structures of singlet, triplet and quintet states are all summarized above. The spin populations of some important atoms are shown in the right corner. The key instances are also labeled in the structures in the unit of Å. The energy of each structure is also given in the figure.

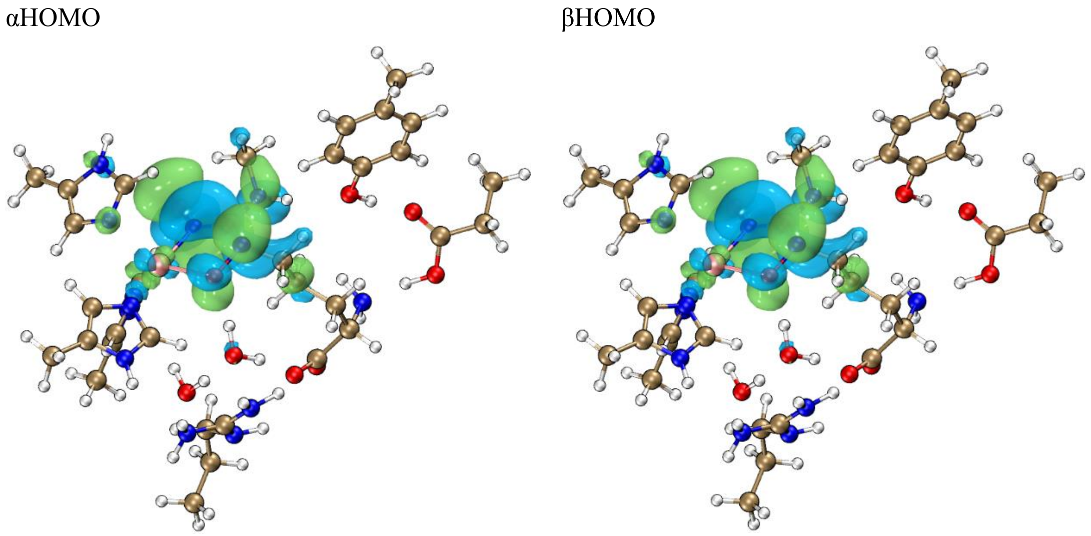


**Figure S3.** Frontier molecular orbitals of $^{3}IntA_{HF}$ without the oxygen binding, in which the HOMOs locate among the $N^{\omega}O$ and $N^{\delta}O$ groups, and $C^{\varepsilon}$ and $N^{\omega}_{Me}$ atoms.

Quintet ($^{5}$IntB)

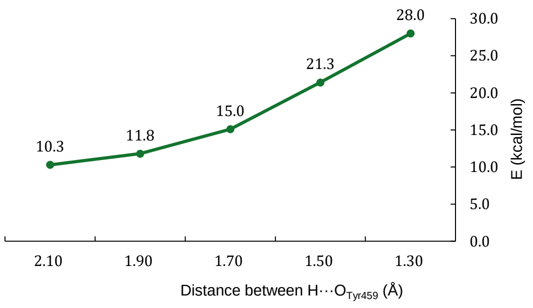


**Figure S4.** The energy variations along the coordination of $N^{\omega}_{Me}$ deprotonation by the Tyr459, which is obtained from the scan calculation based on $^{5}$IntB with the step size of 0.2 Å.

(a) Triplet ($^{3}$IntC) (b) Quintet ($^{5}$IntC)

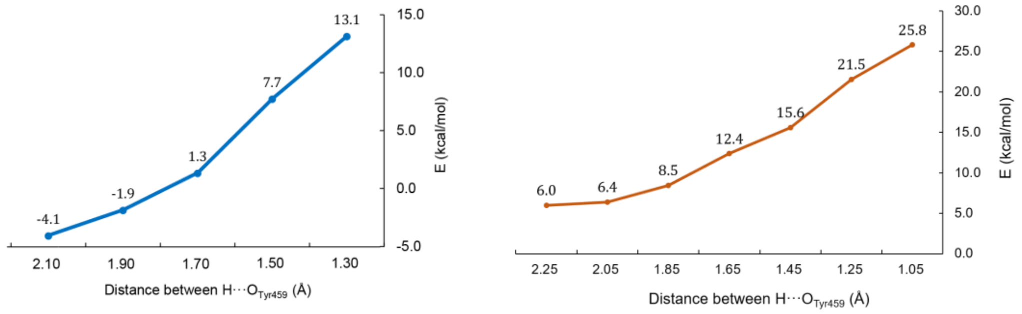


**Figure S5.** The scan calculation along the coordination of $N^{\omega}_{Me}$ deprotonation by the Tyr459 with 0.2 Å step size. (a) shows the energy changes along the proton transfer step from $^{3}$IntC and the energy variations in figure (b) is obtained based on $^{5}$IntC.

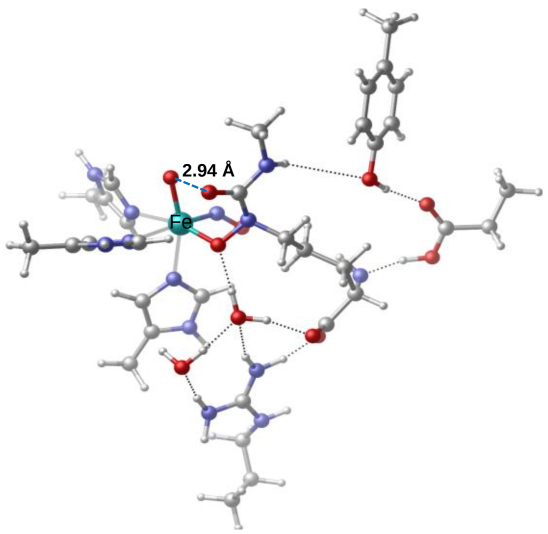


**Figure S6.** The minimum energy crossing point (MECP) between $^{3}$TS_CD and $^{5}$IntD. The MECP was optimized by sobMECP procedure. The energy of this MECP is -31.0 kcal/mol.

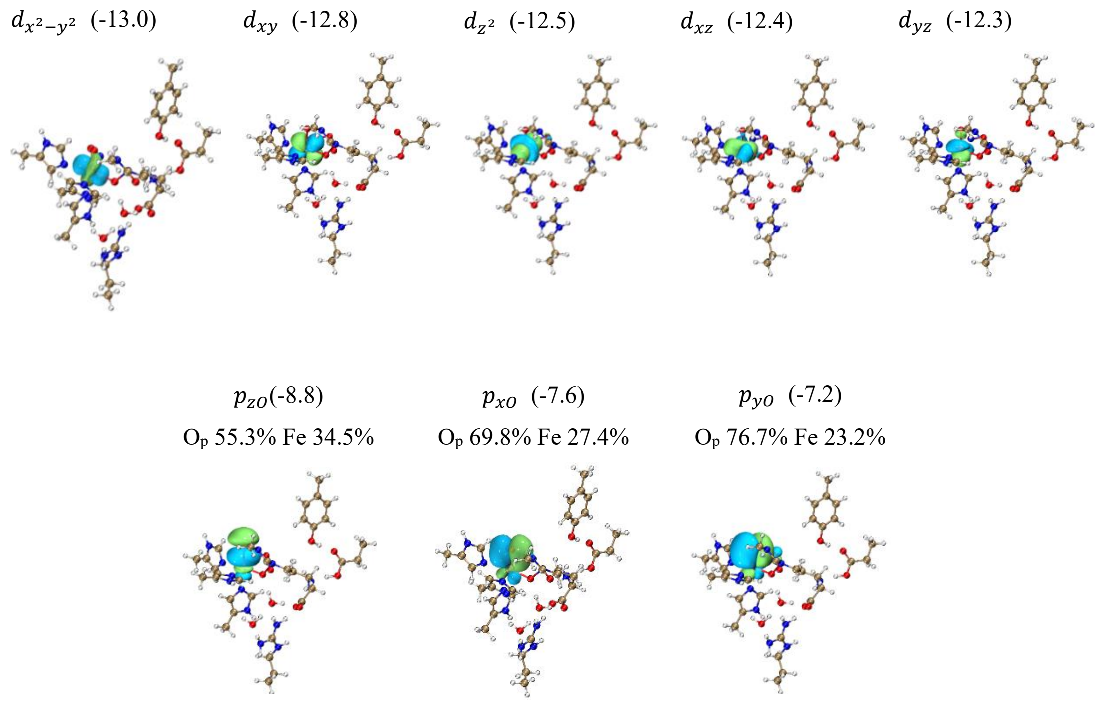


**Figure S7.** The Pipek-Mezey localized molecular orbitals (LMOs) analyses of $Fe^{III}$=O group in $^{5}$IntD. The LMOs on the top are α *d*-orbitals of Fe atom, and the LMOs on the bottom are β *p*-orbitals of O atoms. The energies are labeled in eV.

### Part II The energy profile and O-O cleavage structures of Path IV.

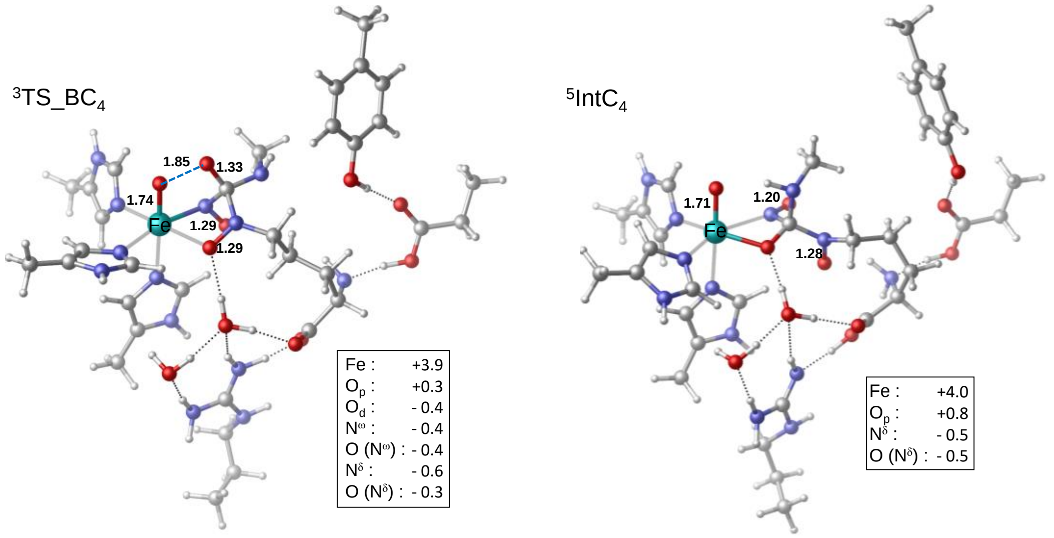


**Figure S8.** The transition structure of O-O homolysis ($^{3}$TS_BC$_{4}$) and the intermediate structure after O-O cleavage ($^{5}$IntC$_{4}$) in Path IV.

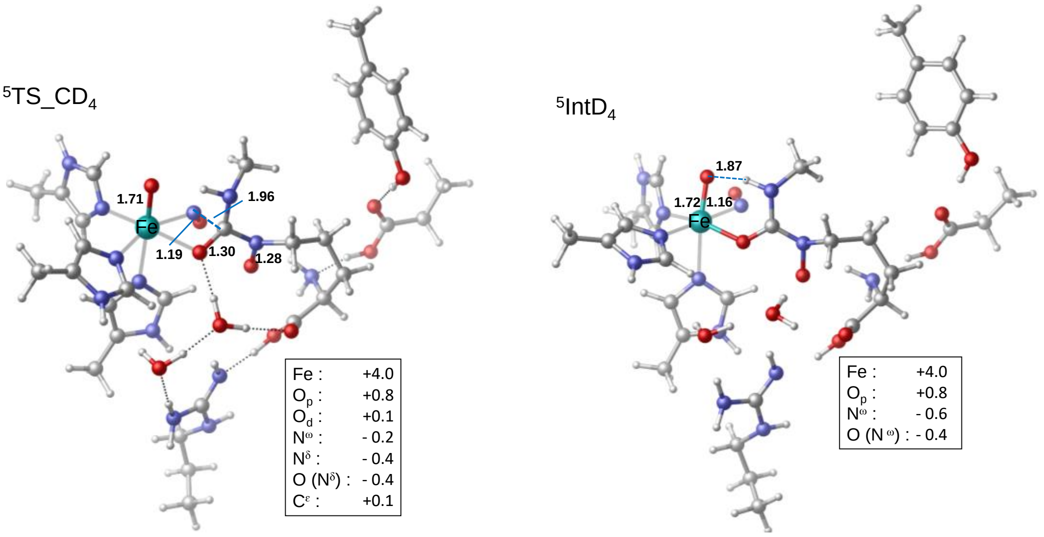


**Figure S9.** The transition structure of C-N rupture ($^{5}$TS_CD$_{4}$) and the intermediate structure after C-N breaking ($^{5}$IntD$_{4}$) in Path IV.

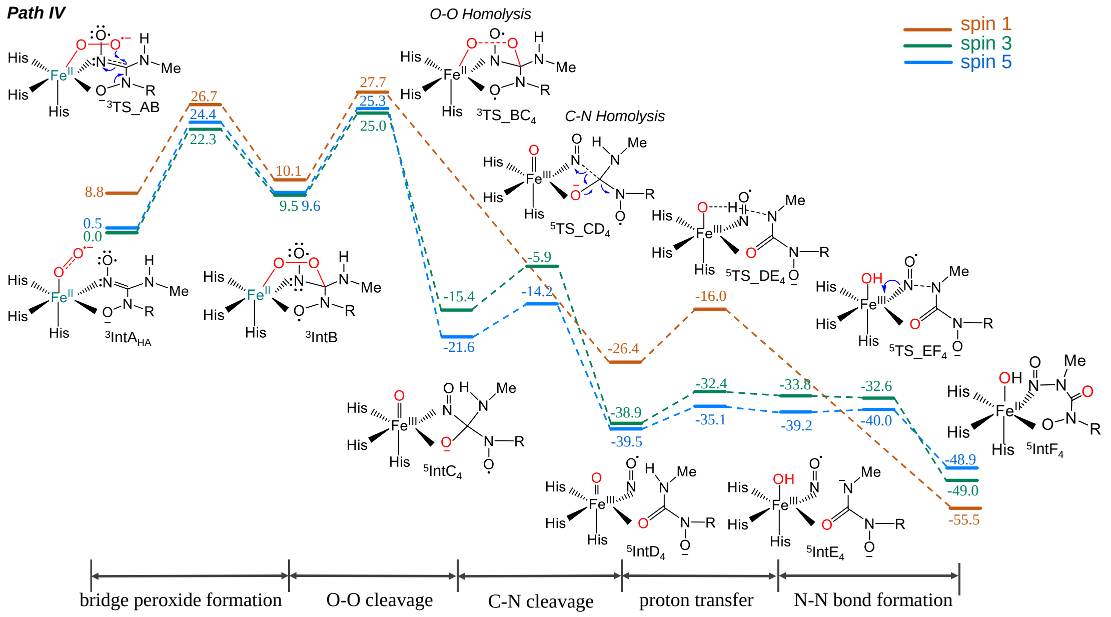


**Figure S10.** Free energy profile of the intramolecular oxidative rearrangement along Path IV. The relative energies are labeled in kcal/mol.

### Part III The energy profile and structure of diaziridine intermediate in Path II.

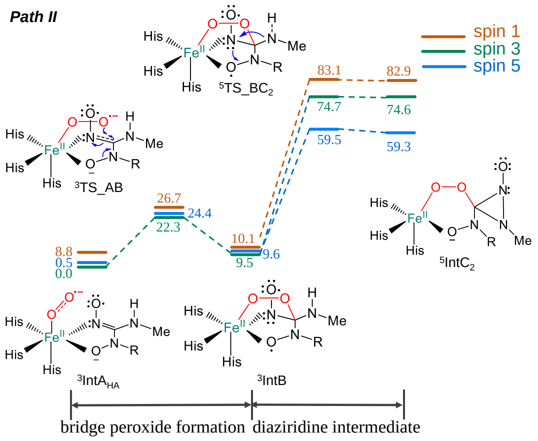


**Figure S11**. The free energy profile of diaziridine intermediate forming steps in Path II.

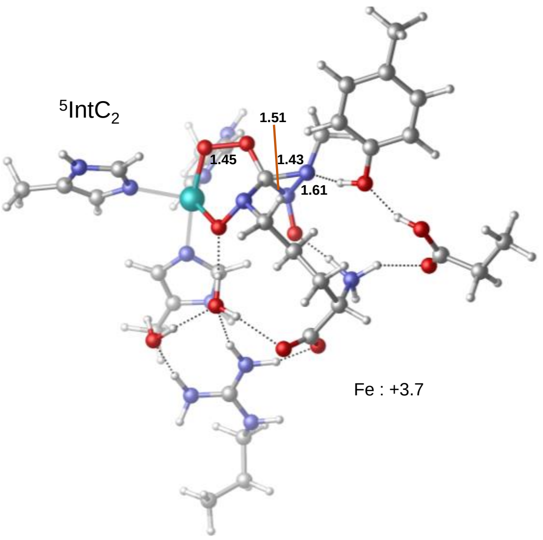


**Figure S12.** Structure of the diaziridine intermediate ($^5$IntC$_2$).
